# Agonism of mosquito and human transient receptor potential ankyrin 1 (TRPA1) channels by the natural drimane sesquiterpene cinnamodial

**DOI:** 10.64898/2025.12.09.693172

**Authors:** Tae Lee, Jayce Liu, Yeaeun Park, H. Liva Rakotondraibe, Xiaolin Cheng, Peter M. Piermarini

## Abstract

TRPA1 channels are multi-modal receptors in animals for sensing noxious physical and chemical factors. They are considered promising biochemical targets for developing repellents of arthropod disease vectors and drugs to treat a variety of medical conditions including pain, inflammation, and itch. Recently, we discovered that cinnamodial (CDIAL), a natural drimane sesquiterpene produced by the medicinal plant *Cinnamosma fragrans*, agonizes mosquito TRPA1 channels and is antifeedant and repellent to *Aedes aegypti*. However, the selectivity of CDIAL for mosquito vs. human TRPA1 and how CDIAL binds to TRPA1 channels are unknown. Here we characterize the agonism by CDIAL of *Aedes aegypti* (Aa) and *Homo sapiens* (Hs) TRPA1 channels and compare it to that induced by two other known TRPA1 agonists, nepetalactone and JT010. Heterologous expression of AaTRPA1 and HsTRPA1 in *Xenopus laevis* oocytes revealed that CDIAL agonized both channels with similar potency. Nepetalactone was less potent of an agonist than CDIAL for both AaTRPA1 and HsTRPA1, but was a more specific agonist for AaTRPA1 over HsTRPA1 compared to CDIAL. JT010 did not detectably activate AaTRPA1, but was a more potent agonist of HsTRPA1 than CDIAL. To generate insights into a putative binding site of CDIAL on AaTRPA1 we mutated six Lys residues (656, 678, 681, 728, 738, 744) and/or a Cys residue (684) in the coupling domain previously hypothesized to be involved. Simultaneous mutation of the six Lys residues to Ala dampened agonistic responses of AaTRPA1 to CDIAL, whereas changing Cys684 to Ser had no detectable impact. Protein modeling and molecular docking simulations for both AaTRPA1 and HsTRPA1 suggest species-specific CDIAL-binding mechanisms to the coupling domain, which involve Lys681 in AaTRPA1 and Cys621 in HsTRPA1.

## 1. Introduction

Transient Receptor Potential Ankyrin 1 (TRPA1) channels are highly-conserved multimodal receptors that are best studied for their roles in sensory neurons of animals where they detect potentially noxious stimuli in the environment, including damaging heat, chemicals (e.g., electrophiles), and radiation (e.g., ultraviolet light) (Talavera et al., 2020). When activated by these stimuli, TRPA1 channels typically stimulate pain and/or avoidance behaviors in animals. However, in some predatory or hematophagous organisms (e.g., pit vipers, ticks, mosquitoes) neuronal TRPA1 is also involved in detecting infrared radiation to facilitate prey/host seeking (Chandel et al., 2024; Gracheva et al., 2010; Mitchell et al., 2017). Beyond neurons, TRPA1 channels are expressed in a wide variety of cells outside of the nervous system including muscle, endothelia, immune cells, and epithelia where they can play roles in sensing molecules associated with cell or tissue damage (e.g., ROS) and bacterial infection (e.g., bacterial lipopolysaccharides) (Talavera et al., 2020). In humans, TRPA1 is considered a potentially valuable target for drug development given its roles in a variety of pathophysiological conditions, including inflammation, itch, and pain (Talavera et al., 2020; Zhang et al., 2023).

In hematophagous adult female mosquitoes, TRPA1 channels play roles in temperature-guided host seeking behaviors, preferred temperature selection, and the avoidance of noxious chemicals (Chandel et al., 2024; Corfas and Vosshall, 2015; Inocente et al., 2018; Li et al., 2019; Lv et al., 2025; Maekawa et al., 2011; Melo et al., 2021; Park and Piermarini, 2025; Wang et al., 2009). Exposure of adult female mosquitoes to chemical agonists of TRPA1 (e.g., allyl isothiocyanate, citronellal, nepetalactone) induces avoidance behaviors and/or disrupts host-seeking (Corfas and Vosshall, 2015; Inocente et al., 2019; Inocente et al., 2018; Kwon et al., 2010; Lv et al., 2025; Maekawa et al., 2011; Melo et al., 2021; Park and Piermarini, 2025). Thus, TRPA1 is considered a valuable biochemical target for development of mosquito repellents (Kang et al., 2010; Salgado, 2017).

We previously demonstrated that cinnamodial (CDIAL)—a natural drimane sesquiterpene dialdehyde produced by medicinal plants of Madagascar (*Cinnamosma* sp.)—agonizes TRPA1 of the malaria mosquito *Anopheles gambiae* (AgTRPA1) when expressed heterologously in *Xenopus* oocytes (Inocente et al., 2018). Moreover, in two wild-type strains (Liverpool & Orlando) of the yellow fever mosquito *Aedes aegypti*, CDIAL was a potent antifeedant and/or repellent; CDIAL showed relatively weak antifeedant activity in a TRPA1 knockout strain of *A. aegypti*, suggesting its mode of action involves modulation of *A. aegypti* TRPA1 (AaTRPA1)(Inocente et al., 2018). Previous studies have not tested whether CDIAL agonizes TRPA1 of mammals, but two other drimane sesquiterpene dialdehydes that are structurally very similar to CDIAL (i.e., polygodial, warburganal) are known agonists of human and mouse TRPA1 (Escalera et al., 2008; Mathie et al., 2017).

The binding mechanisms of sesquiterpene dialdehydes to TRPA1 channels are not well understood, but appear unique compared to unsaturated aldehydes (e.g., acrolein, cinnamaldehyde) or isothiocyanates (e.g., allyl isothiocyanate), which interact primarily with one or more cysteine residues in the coupling/linker domain of the large cytosolic NH_2_-terminus of TRPA1 (Hinman et al., 2006; Kang et al., 2010; Macpherson et al., 2007). However, these cysteine residues do not coordinate binding of sesquiterpene dialdehydes, such as polygodial and isovelleral (Escalera et al., 2008), which are more reactive towards lysine than cysteine (Mathie et al., 2017). Using molecular docking simulations of CDIAL with a predicted structural model of AgTRPA1, we previously identified several lysine residues in the coupling domain (Lys656, Lys678, Lys728, Lys738, and Lys744) that are potentially involved with the binding of CDIAL along with a cysteine residue (Cys684) (Manwill et al., 2020). All of these residues are conserved in AaTRPA1, whereas only three are conserved in the respective coupling domain of *Homo sapiens* TRPA1 (HsTRPA1; Fig. S1). Thus, CDIAL may preferentially agonize AaTRPA1 over HsTRPA1. However, the respective agonistic activities of CDIAL against AaTRPA1 and HsTRPA1 have not previously been determined. Moreover, direct experimental evidence for roles of the above Lys and/or Cys residues in the binding of CDIAL to mosquito TRPA1 channels is lacking.

To start addressing these gaps of knowledge, the first goal of the present study was to characterize the agonistic activity of CDIAL against both AaTRPA1 and HsTRPA1. We hypothesized that CDIAL was a more potent agonist of AaTRPA1 vs. HsTRPA1 given the higher conservation of Lys residues previously predicted to interact with AgTRPA1 (Manwill et al., 2020). Furthermore, we compared the agonistic activity of CDIAL to that of nepetalactone and JT010. Nepetalactone is a highly specific natural agonist of insect TRPA1 channels, including AaTRPA1, that nominally agonizes HsTRPA1 (Li et al., 2025; Melo et al., 2021). JT010 is a highly specific, synthetic agonist of HsTRPA1 (Matsubara et al., 2022; Takaya et al., 2015) that has not previously been tested on insect TRPA1 channels. The second goal of our study was to test the hypothesis that the aforementioned Lys and Cys residues of AaTRPA1 (Fig. S1) contribute to its interaction with CDIAL.

## 2. Materials and Methods

### 2.1. Chemical agonists

CDIAL was extracted from the root bark of *Cinnamosma fragrans* as previously described (Inocente et al., 2019; Inocente et al., 2018; Manwill et al., 2020). In short, air-dried bark was pulverized into powders and extracted with dichloromethane for 5 days at room temperature. Using column chromatography over silica gel, *C. fragrans* extracts were divided into fractions with a gradient of hexanes-ethyl acetate. Fractionated CDIAL was recrystallized using hexanes-ethyl acetate. Nepetalactone and JT010 were purchased from MedChemExpress (Monmouth Junction, NJ), whereas flufenamic acid (FFA) was purchased from Acros Organics (Thermo Scientific, Waltham, MA).

CDIAL and JT010 were each dissolved in DMSO (Thermo Fisher Scientific) as 0.1, 1.0, and 10 mM stock solutions. Nepetalactone was dissolved in DMSO as 10, 100, and 1000 mM stock solutions. FFA was dissolved in DMSO as a 100 mM stock solution. For experiments, stock solutions were diluted 1000-fold in ND96 solution (0.1% DMSO). The ND96 consisted of 96 mM NaCl, 2 mM KCl, 1.8 mM CaCl_2_, 1.0 mM MgCl_2_, and 5 mM HEPES (pH 7.5 titrated with 1 M *N*-methyl D-glucamine; final osmolality = 195 ± 5 mOsmol/kg) (Park and Piermarini, 2025).

### 2.2. AaTRPA1 and HsTRPA1 cDNA constructs

We used a previously developed cDNA construct for AaTRPA1 (Park and Piermarini, 2025), which corresponds to AaTRPA1C (GenBank Accession #LC438794.1) in Li et al. (Li et al., 2019). Genewiz (Azenta Life Sciences, South Plainfield, NJ) was hired to conduct mutagenesis on the AaTRPA1 cDNA to change cysteine 684 into a serine and/or multiple lysine residues (656, 678, 681, 728, 738, 744) into alanine residues. A total of 3 mutant AaTRPA1 cDNA constructs were generated: 1) Cys684Ser; 2) Lys656, 678, 681, 728, 738, 744Ala (Lys_null); and 3) Cys684Ser + Lys_null. The HsTRPA1 cDNA construct was generously provided by Drs. Shigeru Saito and Makoto Tominaga (National Institute for Physiological Science, Okazaki, Japan) (Gupta et al., 2016) and corresponds to GenBank Accession #XP_011515926.

### 2.3. Heterologous expression in Xenopus laevis oocytes

AaTRPA1 cDNA plasmids (wild-type or mutant) were linearized using NotI restriction enzyme (Thermo Fisher, Waltham, MA). The HsTRPA1 cDNA plasmid was linearized using MluI restriction enzyme (Thermo Fisher). The linearized plasmids were used as templates to synthesize complementary RNA (cRNA) using a mMessage mMachine T7 Transcription kit (Invitrogen, Thermo Fisher Scientific, Waltham, MA) for the AaTRPA1 constructs or a mMessage mMachine SP6 Transcription kit (Invitrogen) for the HsTRPA1 construct. All cRNA samples were aliquoted and stored at -80 °C until use.

Each cRNA was injected into defolliculated *Xenopus laevis* oocytes (Ecocyte Bioscience, Austin, TX) for heterologous expression. The oocytes were either injected with 25.2 ng of cRNA (0.9 ng/nl, 28 nl total) or nuclease-free H_2_O (28 nl) for controls. Prior to use in experiments, the injected oocytes were cultured in sterile OR3 media for 2-3 days at 18°C to allow for translation of TRPA1 cRNA into protein. The OR3 media consisted of Leibovitz L-15 media with L-glutamine (Gibco,Thermo Fisher Scientific), 1000 U penicillin-streptomycin (Gibco,Thermo Fisher Scientific), and 5 mM HEPES (pH 7.5 titrated with 1 N NaOH; final osmolality = 195 ± 5 mOsmol/kg) (Romero et al., 1998).

### 2.4. Two-electrode voltage clamping (TEVC) of Xenopus oocytes

TEVC experiments were performed using an established protocol for measuring TRPA1 agonism (Inocente et al., 2018; Park and Piermarini, 2025). In brief, an oocyte was placed in a RC-3Z chamber (Warner Instruments, Holliston, MA) with gravity-fed ND96 solution (∼3 ml/min). The oocyte was then impaled with two glass electrodes (each filled with 3 M KCl; 0.5-1 MΩ tip resistance) to respectively measure membrane potential (V_m_) and whole-cell membrane current (I_m_). The V_m_ and I_m_ readings were measured and recorded with an OC-725 oocyte clamp (Warner Instruments) bridged to pCLAMP software (Axoscope, Version 10.7, Molecular Devices) via a MiniDigi-1A interface (Molecular Devices). Upon stabilization of V_m_ (∼2 min), the oocyte was clamped to a hyperpolarizing potential of 30 mV relative to the resting V_m_ to promote inward (negative) membrane currents when TRPA1 was activated (Inocente et al., 2018; Park and Piermarini, 2025).

Once the resting I_m_ stabilized (∼2 min), the oocyte bath solution was changed to ND96 containing a chemical agonist using an 8-way rotary valve (Model 5012, Rheodyne, Ronhert Park, CA). After 40 s, the bath solution was switched back to ND96 for approximately 2 min to wash out the agonist before applying the next agonist or concentration. For concentration-response experiments, three concentrations of CDIAL (0.1 μM, 1 μM, 10 μM), nepetalactone (10 μM, 100 μM, 1000 μM), or JT010 (0.1 μM, 1 μM, 10 μM) were applied consecutively to the same oocyte starting from the lowest to highest concentration with washes of ND96 between each concentration. In experiments with nepetalactone, each oocyte was treated with 10 μM CDIAL after the last nepetalactone treatment (1000 μM); in this context, CDIAL served as a reference compound to compare the relative responses of 1000 μM netpetalactone to 10 μM CDIAL. For controls, H_2_O-injected oocytes were exposed to the same concentrations and orders of chemical agonists. H_2_O-injected oocytes showed no detectable responses to any of the agonists (Fig. S2).

The agonistic response of an oocyte to a particular chemical or concentration of that chemical was defined as the maximum change in I_m_ (ΔI_m_) elicited (Park and Piermarini, 2025). The ΔI_m_ was measured using Clampfit 10.7 (Molecular Devices). For concentration-response experiments, each ΔI_m_ value was standardized to that for the highest concentration of the respective agonist. EC_50_ values for agonists were estimated based on fitting a non-linear curve to the standardized data using the ‘log(agonist) vs. normalized response’ method in Prism (v.10.6, Graphpad Software Inc., La Jolla, CA). If an agonist only induced a detectable response at the highest concentration tested then a curve was not fitted and an EC_50_ was not estimated. For agonists that yielded an EC_50_ for both AaTRPA1 and HsTRPA1, the respective EC_50_ values were compared using an extra sum of squares F-test (α=0.05) in Prism. In some cases, the ΔI_m_ elicited by 1000 μM nepetalactone was compared to that of 10 μM CDIAL using a Mann-Whitney rank sum test (α=0.05) in Prism (data were not normally distributed as determined by a Shapiro-Wilk test).

### 2.5. Effects of Cys and/or Lys mutations on binding of CDIAL to AaTRPA1

To compare agonism by CDIAL among the wild-type (WT) and mutant AaTRPA1 constructs, each oocyte was first treated with FFA (100 μM) for 40 s as a reference agonist to account for potential differences in heterologous expression among the constructs (Hu et al., 2009; Wang et al., 2012). The oocyte was then rinsed with ND96 for approximately 2 min to wash out the FFA and allow I_m_ to return to the baseline. The oocyte was then treated with 1 μM CDIAL for 40 s.

The ΔI_m_ was measured for both FFA and CDIAL using Clampfit 10.7 as described above. For comparisons among the WT and mutants, the ΔI_m_ for CDIAL was normalized to that for FFA, resulting in a CDIAL:FFA ratio. To avoid skewed CDIAL:FFA ratios resulting from weak TRPA1 expression, we excluded oocytes with ΔI_m_ responses of less than 0.1 µA for FFA. The CDIAL:FFA ratios of the WT and mutant strains were compared using a non-parametric one-way ANOVA (Kruskal-Wallis test with multiple comparisons made using Dunn’s Method, α=0.05) in Prism (data were not normally distributed as determined by a Shapiro-Wilk test).

### 2.6. AaTRPA1-CDIAL complex structure prediction

The structure of AaTRPA1 (UniProt A0A679BVR2) in complex with CDIAL (PubChem CID 442354) was modeled using AlphaFold 3 (AF3)(Abramson et al., 2024). To reduce computational cost without compromising the structural features relevant to ligand binding, only residues 500–1246 of AaTRPA1 were included. A two-step docking strategy was considered: 1) non-covalent docking to identify the potential conjugation residue and covalent attachment site of CDIAL; and 2) covalent docking to model the final conjugated complex.

#### 2.6.a. Noncovalent docking

AF3 predictions were performed with 200 random seeds, each generating 5 models (1,000 total). Poses were retained if CDIAL was localized near the proposed lysine pocket (Lys656/Lys678/Lys681/Lys728/Lys738/Lys744) and exhibited no severe steric clashes based on Maestro Contacts (default settings).

#### 2.6.b. Covalent docking

The terminal nitrogen atom (Nε) of Lys681 was designated as the covalent attachment site. Three carbon atoms on CDIAL (C7, C11, and C12) were tested for covalent linkage (Fig. 1). The covalent bond was defined between Lys Nε and the designated ligand carbon. To mimic imine dehydration, oxygen atoms were removed for C11- and C12-linked CDIAL variants using AF3’s SMILES-to-cif conversion tool; the C7-linked variant was left unmodified. For each linkage type, 19 random seeds × 5 models were generated (285 total poses).

**Fig. 1.**
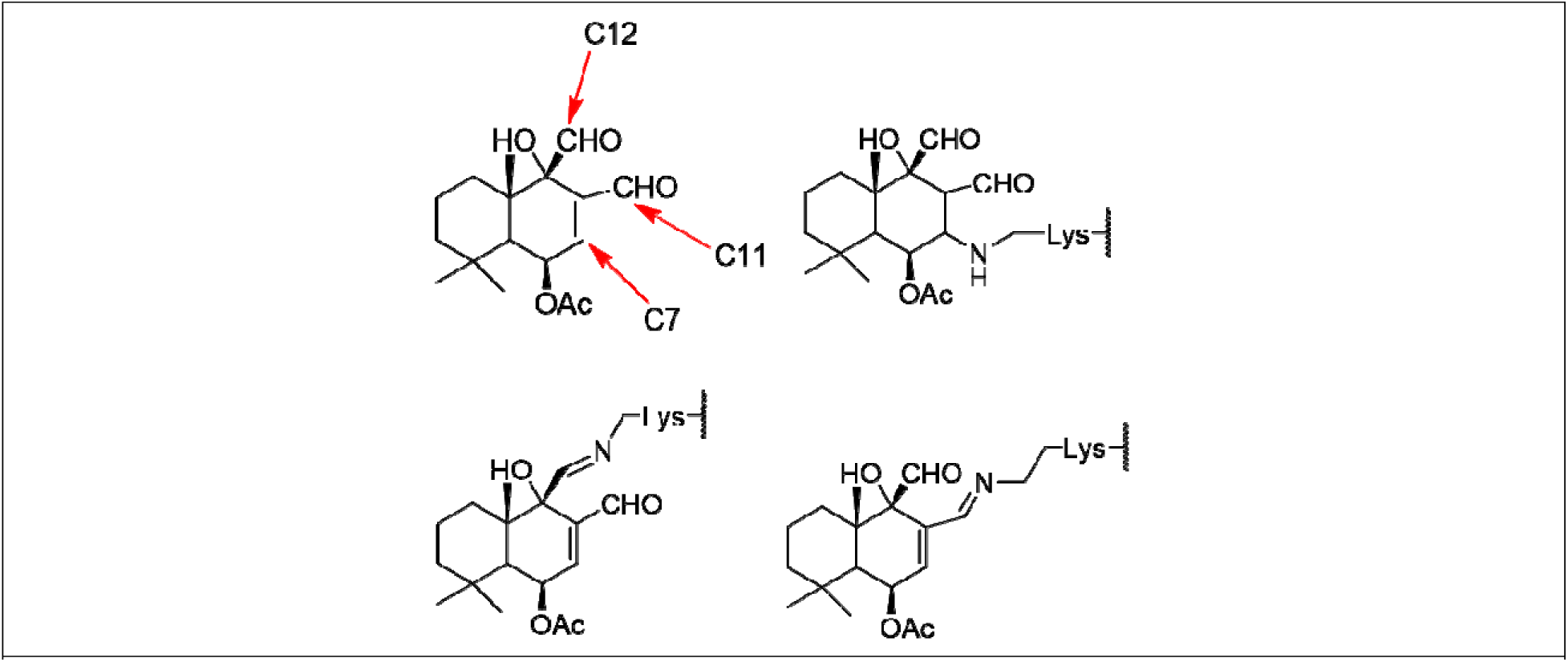
Atoms on CDIAL (C7/C11/C12) used to form covalent bond with the receptor in covalent docking simulations. Arrows indicate the three sites tested. Imine formation was modeled at C11/C12.

All AF3-predicted complexes were hydrogenated and energy-minimized using the Schrödinger Protein Preparation Wizard with default settings (Sastry et al., 2013; Schrödinger, Inc, 2025). Poses were assessed using: 1) steric clash filtering via Maestro Contacts (default settings), 2) root-mean-square deviation (RMSD) analysis of ligand poses (after aligning protein backbone atoms, RMSD was calculated for ligand heavy-atoms; poses with lower RMSDs relative to the noncovalent docking pose were prioritized), 3) local contact analysis where preference was given to poses with enhanced interactions (H-bonding and hydrophobic contact) with pocket residues.

### 2.7. HsTRPA1-CDIAL complex structure prediction

The HsTRPA1 (UniProt O75762, residues 401–1119)-CDIAL complex was modeled using the same AF3 workflow described above.

#### 2.7.a. Noncovalent docking

200 random seeds × 5 models each, yielded 1,000 structures.

#### 2.7.b. Covalent docking

Cys621 (Sγ atom) was selected as the conjugation site, forming a single S–C bond with C7, C11, or C12 of CDIAL to mimic hemithioacetal formation. For each linkage type, 20 seeds × 5 models were generated (300 total). Pose assessment criteria were identical to the AaTRPA1 analysis.

### 2.8. AaTRPA1-JT010 complex structure prediction

The same AaTRPA1 fragment (residues 500–1246) was modeled in complex with JT010 (PubChem CID 18524489) using AF3 without covalent constraints (200 seeds × 5 models; 1,000 total). Models were selected based on proximity of JT010 to the known active site and the proposed CDIAL binding pocket.

## 3. Results

### 3.1. Concentration-response relationships of AaTRPA1 and HsTRPA1 to chemical agonists

In oocytes expressing either AaTRPA1 or HsTRPA1, treatment with 0.1-10 µM CDIAL elicited concentration-dependent agonism (ΔI_m,_ Fig. 2A-B). Fig. 2C shows the mean concentration-response curves to CDIAL for both channels, revealing similar (p = 0.99) estimated EC_50_ values of 1.93 μM (95% C.I. = 1.535 to 2.44 μM) for AaTRPA1 and 2.25 μM (95% C.I. = 1.77 to 2.87 μM) for HsTRPA1.

**Fig. 2.**
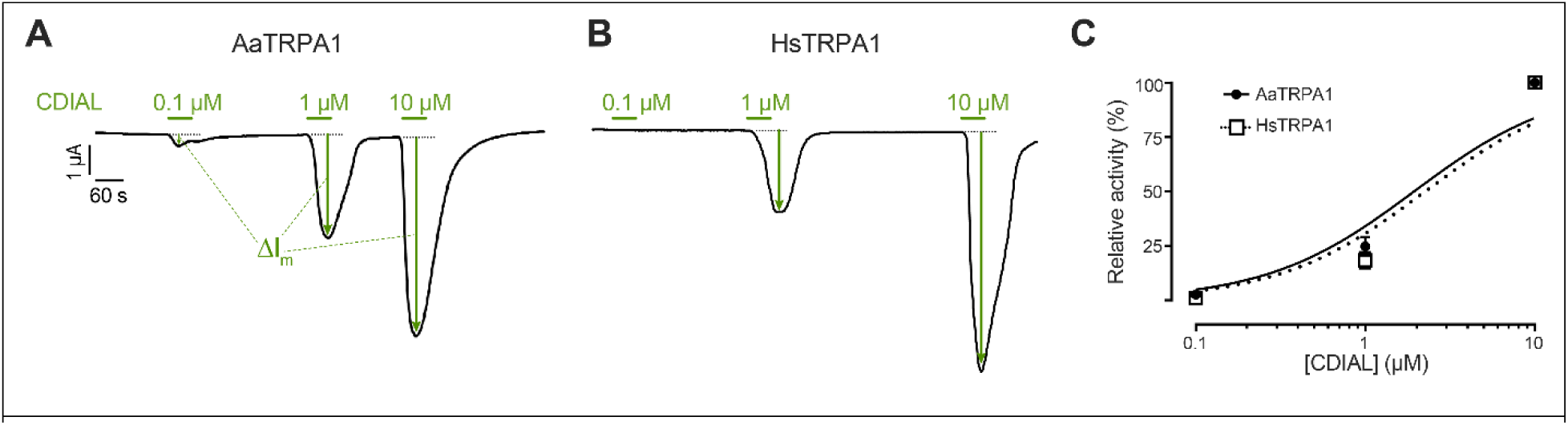
Agonistic effects of CDIAL on AaTRPA1 and HsTRPA1 oocytes. Representative traces of I_m_ in voltage-clamped oocytes expressing AaTRPA1 (A) or HsTRPA1 (B) in response to increasing concentrations of CDIAL. Green horizontal bars indicate exposure of oocytes to CDIAL at the indicated concentrations. Arrows indicate ΔI_m_. C) Concentration-response relationships of CDIAL for AaTRPA1 and HsTRPA1. Values are means ± SEM, based on 19 oocytes for AaTRPA1 and 18 oocytes for HsTRPA1.

Experiments with nepetalactone required higher concentrations (10-1000 µM) to induce concentration-dependent agonism in AaTRPA1 oocytes (Fig. 3A). Subsequent exposure of the same oocytes to 10 µM CDIAL revealed that the degree of AaTRPA1 agonism induced by 1000 µM nepetalactone was relatively weak compared to that induced CDIAL (Fig. 3A). On average, the ΔI_m_ induced by 1000 µM nepetalactone in AaTRPA1 oocytes (n = 11) was 60.2 ± 8.7% lower than that induced by 10 µM CDIAL. In HsTRPA1 oocytes, detectable agonism by nepetalactone was only observed at 1000 µM (Fig. 3B). Subsequent exposure of the same oocytes to 10 µM CDIAL revealed that the degree of HsTRPA1 agonism induced by 1000 µM nepetalactone was dramatically weaker compared to that induced CDIAL (Fig. 3B). On average, the ΔI_m_ induced by 1000 µM nepetalactone in HsTRPA1 oocytes (n = 14) was 98.8 ± 0.46% lower than that induced by 10 µM CDIAL. Fig. 3C shows the concentration-response curve for nepetalactone in AaTRPA1 oocytes, revealing an estimated EC_50_ value of 197.0 μM (95% C.I. = 147.9 to 262.3 μM). The limited responses of HsTRPA1 oocytes to nepetalactone precluded a concentration-response curve and EC_50_ estimate.

**Fig. 3.**
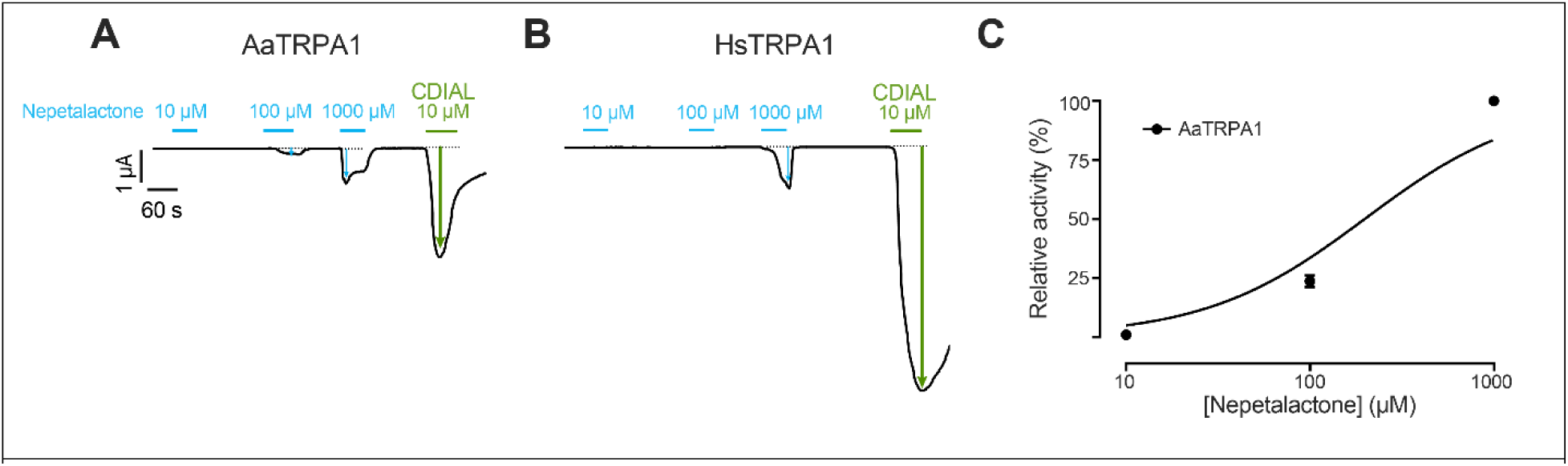
Agonistic effects of nepetalactone on AaTRPA1 and HsTRPA1 oocytes. Representative traces of I_m_ in voltage-clamped oocytes expressing AaTRPA1 (A) or HsTRPA1 (B) in response to increasing concentrations of nepetalactone. Cyan and green horizontal bars indicate exposure of oocytes to nepetalactone and CDIAL, respectively at the indicated concentrations. Arrows indicate ΔI_m_. C) Concentration-response relationship of nepetalactone against AaTRPA1. Values are means ± SEM, based on 11 oocytes for AaTRPA1.

Experiments with 0.1 to 10 µM JT010 did not reveal detectable agonism in AaTRPA1 oocytes (Fig. 4A; n = 5). In contrast, JT010 elicited potent concentration-dependent agonism from 0.1 to 10 µM in HsTRPA1 oocytes (Fig. 4B). Although the AaTRPA1 and HsTRPA1 oocytes exposed to JT010 were not subsequently treated with CDIAL (as occurred in the nepetalactone experiments), independent oocytes within the same batch for both AaTRPA1 and HsTRPA1 showed stronger or similar agonistic responses, respectively, to 10 µM CDIAL compared to those induced by 10 µM JT010 (Fig. S3). Fig. 4C shows the concentration-response curve for JT010 in HsTRPA1 oocytes, revealing an estimated EC_50_ value of 0.80 μM (95% C.I. = 0.49 to 1.1 μM). The lack of detectable responses of AaTRPA1 oocytes to JT010 precluded a concentration-response curve and EC_50_ estimate.

**Fig. 4.**
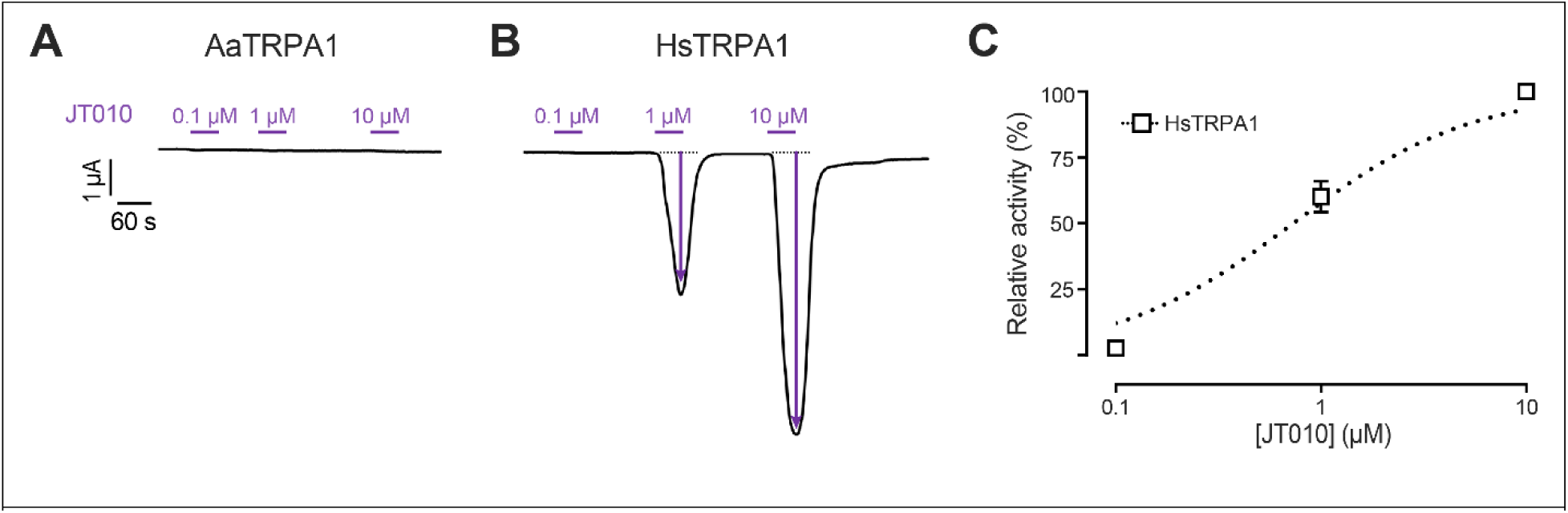
Agonistic effects of JT010 on AaTRPA1 and HsTRPA1 oocytes. Representative traces of I_m_ in voltage-clamped oocytes expressing AaTRPA1 (A) and HsTRPA1 (B) in response to increasing concentrations of JT010. Purple horizontal bars indicate exposure of oocytes to JT010 at the indicated concentrations. Arrows indicate ΔI_m_. C) Concentration-response relationship of JT010 against HsTRPA1. Values are means ± SEM, based on 4 oocytes for HsTRPA1.

### 3.2. Elucidation of putative CDIAL binding region in AaTRPA1

Previous molecular dynamic simulations by our group (Manwill et al., 2020) led to the hypothesis that CDIAL interacted with Cys684 and/or several Lys residues within the coupling domain of mosquito TRPA1 channels (Fig. S1). To test this hypothesis, we heterologously expressed 3 mutant constructs of AaTRPA1 in which Cys684 was changed to a Ser (Cys684Ser) and/or the six Lys residues were simultaneously changed to Ala (Lys_null). We then compared the relative agonistic responses of these mutants to 1 µM CDIAL with that of wild-type (WT) AaTRPA1. To account for differential CDIAL responses due to potentially variable heterologous expression among the mutants, each oocyte was exposed to 100 µM FFA, a non-electrophilic and reversible agonist of TRPA1 channels (Hu et al., 2010), before treatment with CDIAL.

Fig. 5A shows representative I_m_ traces of voltage-clamped oocytes expressing WT or mutant AaTRPA1. In oocytes expressing WT or Cys684Ser AaTRPA1, 1 µM CDIAL elicited ΔI_m_ responses that were stronger than those of 100 µM FFA. However, in oocytes expressing Lys_null or Cys684Ser + Lys_null AaTRPA1, 1 µM CDIAL elicited ΔI_m_ responses that were weaker than those of 100 µM FFA. Fig. 5B compares the average CDIAL ΔI_m_ responses normalized to those of FFA (i.e., CDIAL/FFA ΔI_m_ ratio) among the WT and mutant AaTRPA1 constructs. Both the WT and Cys684Ser AaTRPA1 had similar CDIAL/FFA ΔI_m_ ratios (WT = 4.91 ± 0.92, Cys684Ser =6.45 ± 1.22; Fig. 5B). Each of these ΔI_m_ ratios was greater than those of Lys_null and Cys684Ser + Lys_null, which were similar to each other (Lys_null = 0.77 ± 0.14, Cys684Ser+Lys_null = 0.54 ± 0.03; Fig. 5B).

**Fig. 5.**
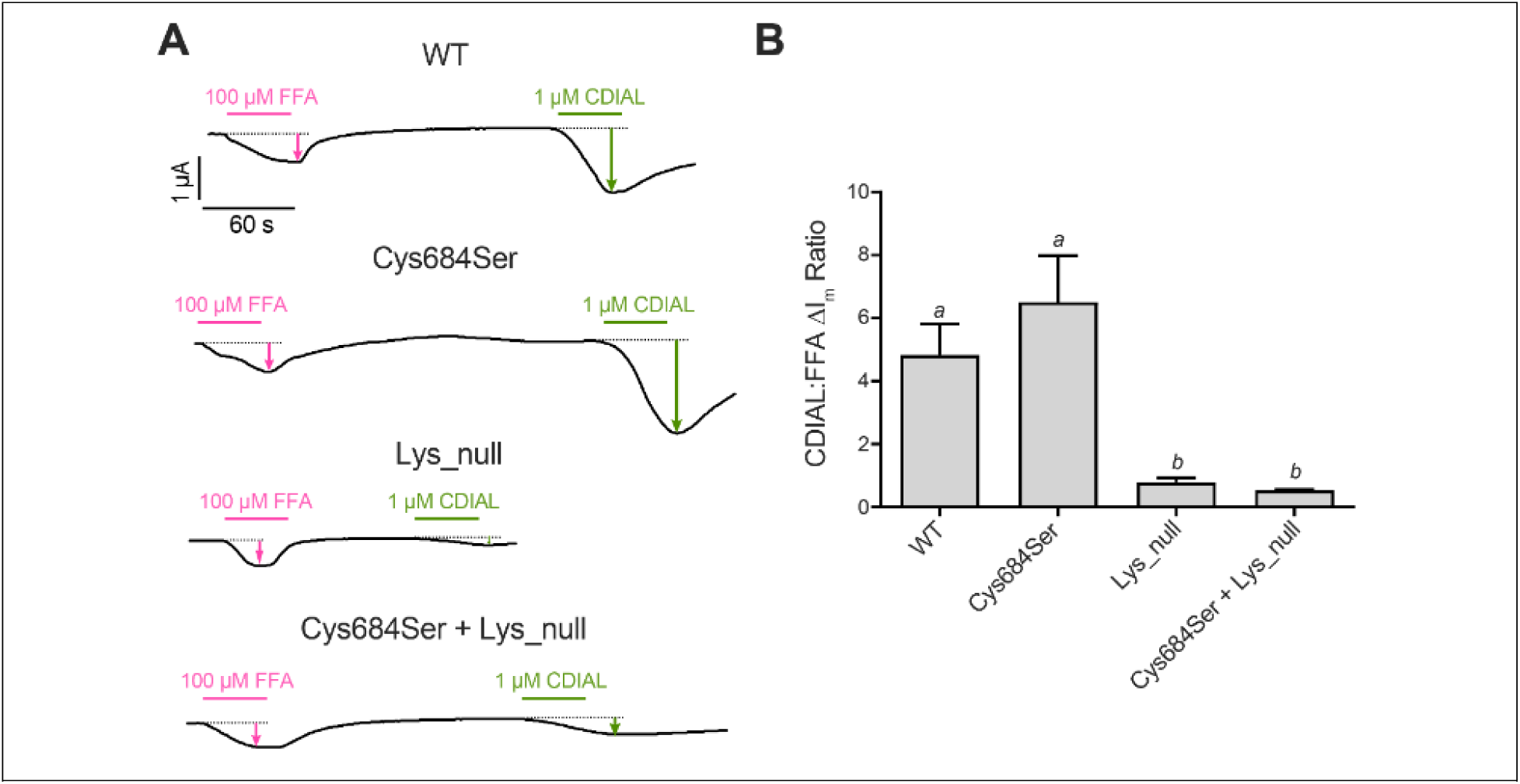
A) Agonistic effects of CDIAL relative to FFA in voltage-clamped oocytes expressing various constructs of AaTRPA1: WT, Cys684Ser, Lys_null, or Cys684Ser+Lys_null. Magenta and green horizontal bars indicate exposure of oocytes to FFA and CDIAL, respectively, at the indicated concentrations. Arrows indicate ΔI_m_. B) Ratios of CDIAL to FFA agonism (ΔI_m_) for each construct. Values are means ± SEM, based on 28, 28, 18, and 23 oocytes respectively for WT, Cys684Ser, Lys_null, and Cys684Ser + Lys_null. Italicized lower-case letters indicate statistical categorization of the means as determined by a Kruskal-Wallis ANOVA with a Dunn’s post-hoc test (P < 0.05).

### 3.3. Structure prediction of the AaTRPA1-CDIAL complex

The mutagenesis results of Fig. 5 suggest that CDIAL binding to AaTRPA1 involves one or more of the six mutated Lys residues and not Cys684. To generate insights into how CDIAL may interact with these Lys residues, we used AlphaFold 3, which can handle covalent binding predictions not easily done with conventional docking approaches. Covalent binding was hypothesized to proceed through a two-step mechanism: first, CDIAL engages in weak but selective noncovalent interactions within the lysine pocket; second, a covalent bond forms between CDIAL and the receptor. This stepwise process may enhance the selectivity of CDIAL binding. One thousand noncovalent AaTRPA1-CDIAL complex structures were predicted. Of these, seven satisfied both the pocket-proximity and clash-free criteria, with all poses placing CDIAL near Lys681 (Fig. 6).

**Fig. 6.**
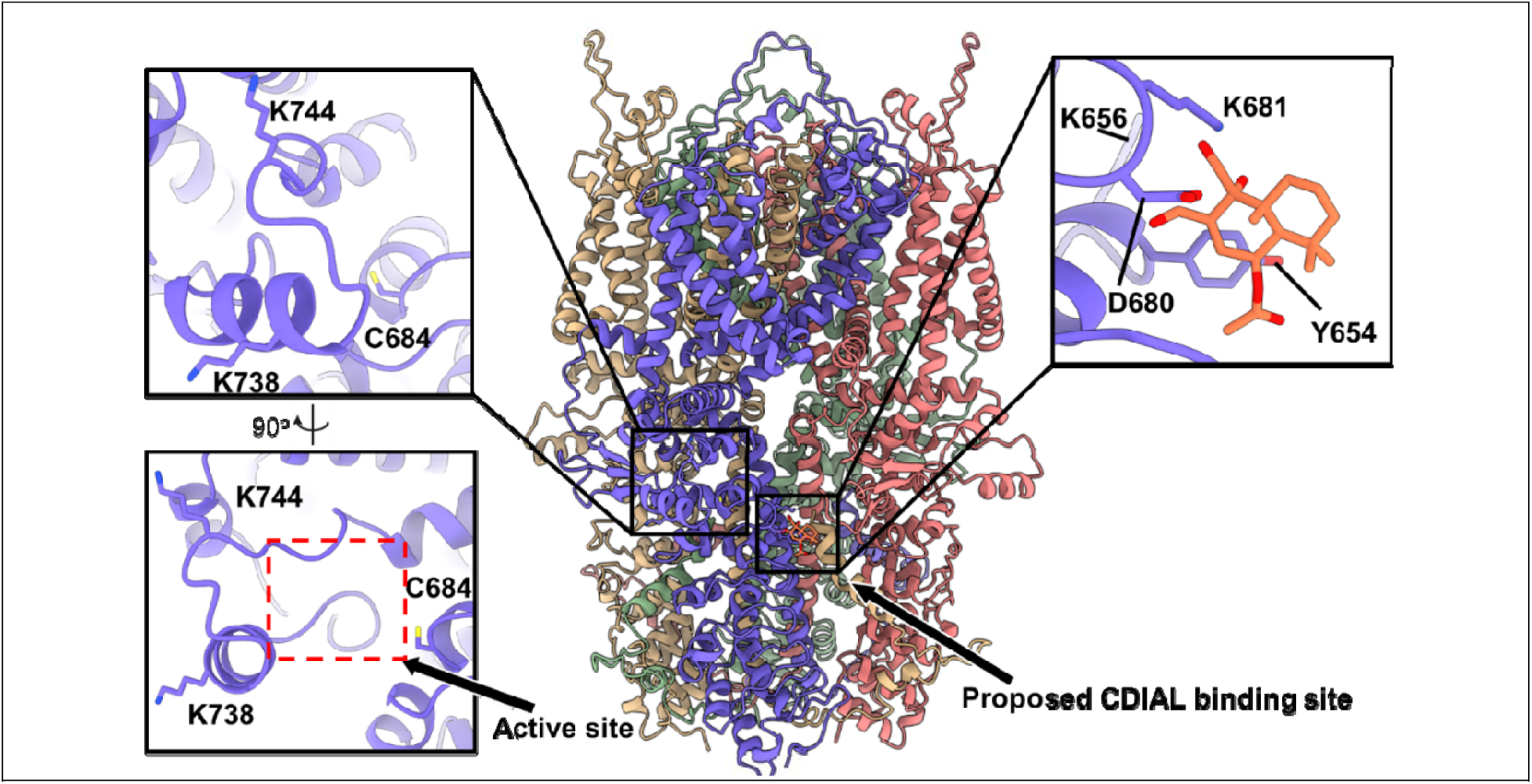
AlphaFold3-predicted AaTRPA1-CDIAL complex. AaTRPA1 is modeled as a homotetramer with chains colored as follows: blue (chain A), pink (chain B), green (chain C), and yellow (chain D). CDIAL is shown in orange. A pocket homologous to the reversible covalent agonist-binding site in HsTRPA1 and the proposed CDIAL-binding region are highlighted and enlarged. Key lysine residues and other residues involved in CDIAL binding are labeled.

Given the absence of cysteine residues in this pocket, we hypothesized that covalent binding could occur through imine formation between Lys681 Nε and one of three possible carbon sites on CDIAL (C7, C11, or C12; Fig. 1). AF3 predictions were generated for each linkage scenario and filtered using the criteria described above. Across all metrics—steric clash rate, covalent-non-covalent pose similarity, and local interaction strength—the C11-linked model emerged as the most favorable (Table 1), showing moderate clash rates, the closest pose clustering, and the highest counts of local hydrogen bonds and hydrophobic contacts.

**Table 1.** Summary of pose analysis for AF3-predicted covalent complexes: AaTRPA1–CDIAL (Lys681) and HsTRPA1–CDIAL (Cys621).

| Series | Carbon atom | Poses inside cluster | Clash rate | Pose that has interaction |
| --- | --- | --- | --- | --- |
| AaTRPA1-CDIAL | C7 | 50/95 | 13/50 (26%) | 4/37 |
|  | C11 | 66/95 | 14/66 (21%) | 20/52 |
|  | C12 | 48/95 | 10/48 (20%) | 1/38 |
| HsTRPA1-CDIAL | C7 | 23/100 | 7/23 (30%) | N/A |
|  | C11 | 32/100 | 7/32 (21%) | N/A |
|  | C12 | 69/100 | 4/59 (7%) | 17/55 |

We then investigated how the binding of CDIAL affects the structure of AaTRPA1 that may be relevant to its potentiation. Compared to apo AaTRPA1 and the noncovalent complex, representative C11-linked models showed new H-bonds and hydrophobic contacts with neighboring residues, including Tyr651, Asp680, Tyr654 (Fig. 6) and they also induced a local rearrangement of the loop harboring Lys681 (RMSD = 6.8 Å) (Fig. S4).

### 3.4. HsTRPA1-CDIAL complex structure prediction

We also generated a model to predict how CDIAL interacts with HsTRPA1 (Fig. 7). Among the 1,000 noncovalent HsTRPA1–CDIAL models, 6 satisfied the active-site proximity and clash-free criteria. None localized to the lysine pocket observed in AaTRPA1, consistent with the absence of such a pocket in HsTRPA1. Instead, all six poses placed CDIAL near Cys621 and reproduced the Pro666 displacement characteristic of agonist engagement, which is also known to occur for JT010 (Fig. 8) (Matsubara et al., 2022). Notably, one pose orientated C7, C11, and C12 favorably toward the Sγ atom of Cys621 (Fig. 8b).

**Fig. 7.**
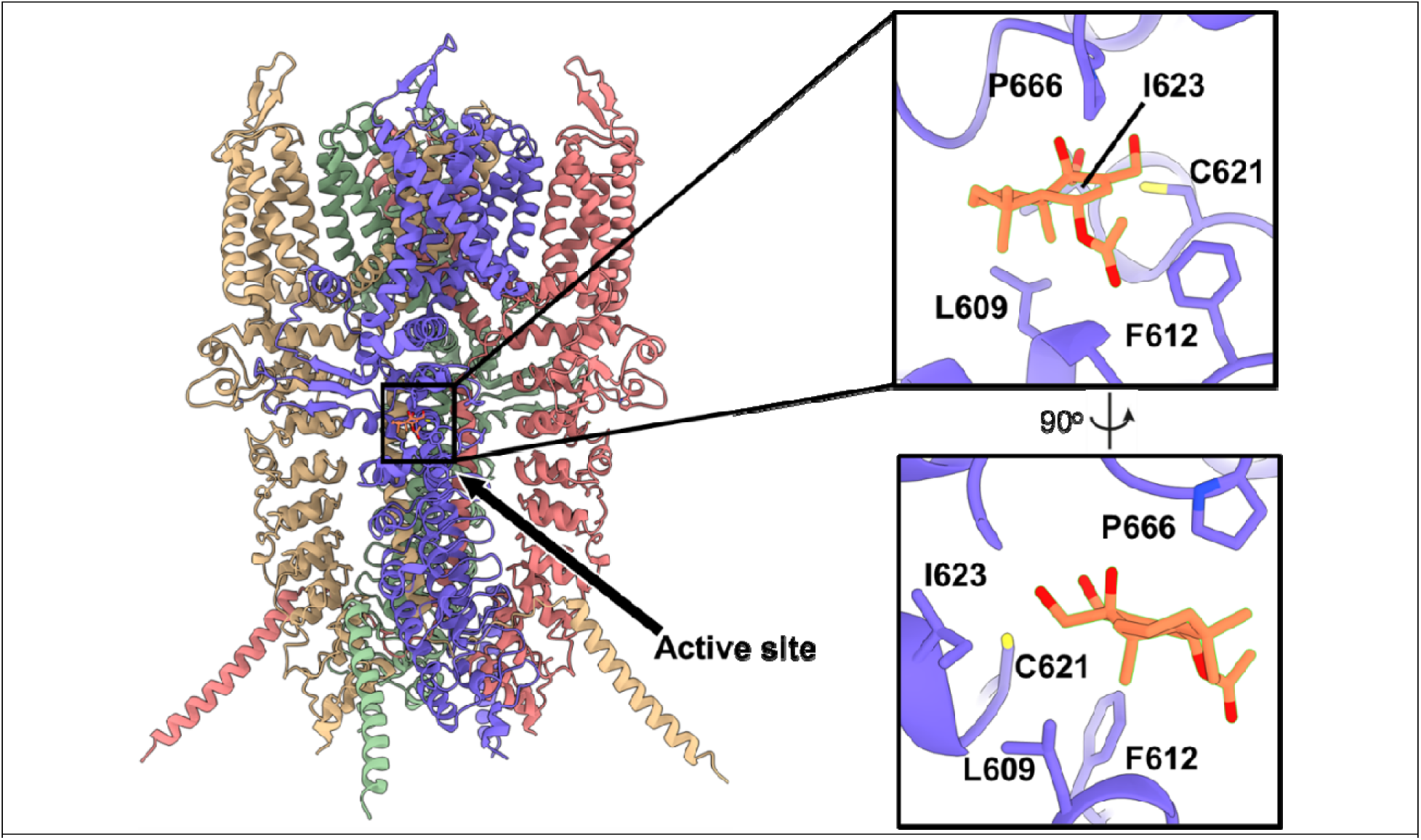
AlphaFold3-predicted HsTRPA1-CDIAL complex. HsTRPA1 is modeled as a homotetramer with chains colored as follows: blue (chain A), pink (chain B), green (chain C), and yellow (chain D). CDIAL is shown in orange. The active site is highlighted and enlarged. Residues contributing to HsTRPA1-CDIAL interaction are labeled.

**Fig. 8.**
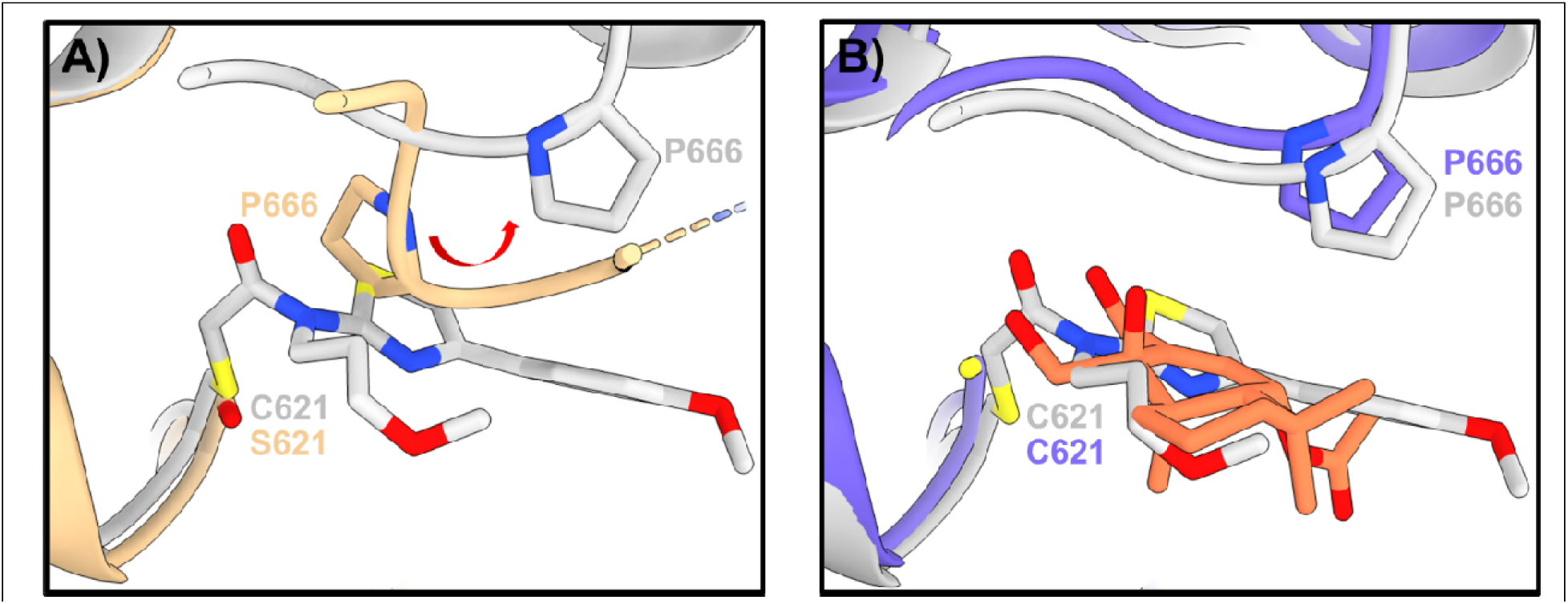
A) Experimental structure showing that covalent binding of JT010 (gray) shifts P666 of HsTRPA1 from the left in apo state (yellow) to the right in the active state. B) Noncovalent binding of CDIAL (blue and orange) to HsTRPA1 caused a similar conformational change of P666 compared to the experimental structure (gray).

### 3.5. AaTRPA1-JT010 complex structure prediction

To gain insight into why JT010 did not agonize AaTRPA1 despite the conserved Cys residue (Cys684) with Cys621 in HsTRPA1, we generated a model to predict how JT010 interacts with AaTRPA1. From 1,000 AF3-predicted noncovalent AaTRPA1–JT010 models, only one satisfied the selection criteria. This pose occupied the active site with the phenyl and thiazole rings oriented toward the pocket rim, enabling π–π and cation–π interactions with edge aromatic residues like His751 and Trp745, while positioning the electrophilic warhead inward to form hydrogen bond with Thr686. This arrangement resembled HsTRPA1-JT010 binding mode. However, the distance between the electrophile and Cys684 remained large (7.7 Å), making covalent bond formation unlikely. Additionally, no conformational changes similar to those in HsTRPA1 upon JT010 binding were observed (Fig. 10).

**Fig. 9.**
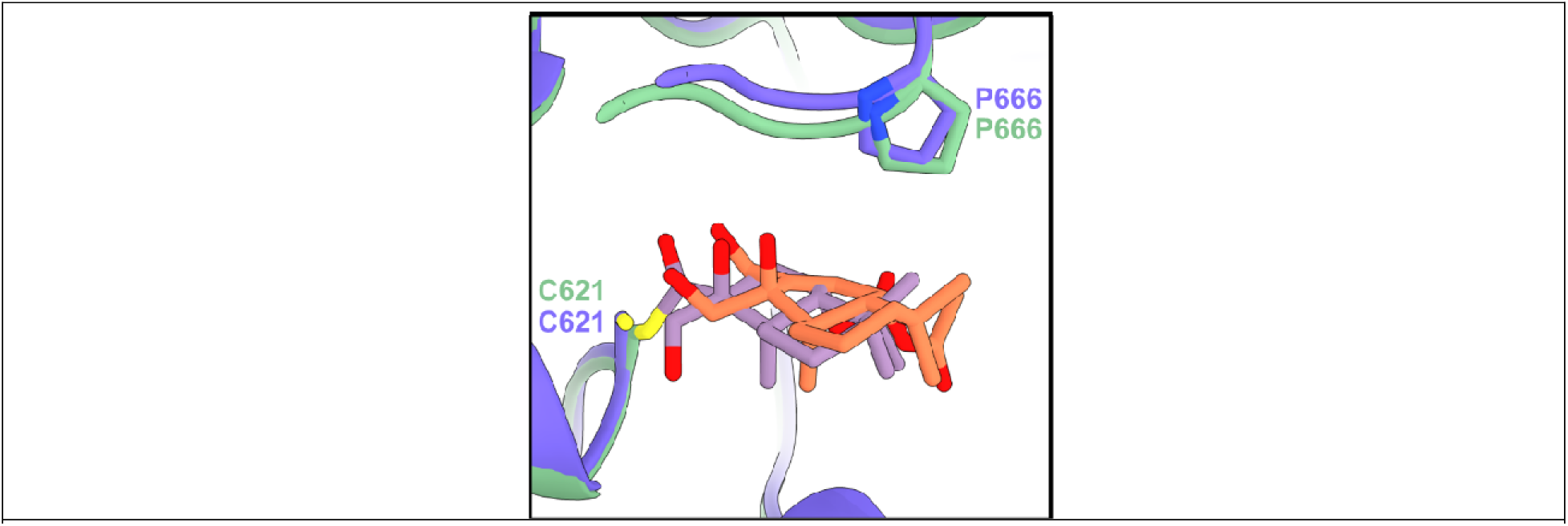
Comparison of covalent and noncovalent binding of CDIAL in the HsTRPA1 active state. Blue and orange represent the noncovalent complex, while green and purple represent the covalent complex.

**Fig. 10.**
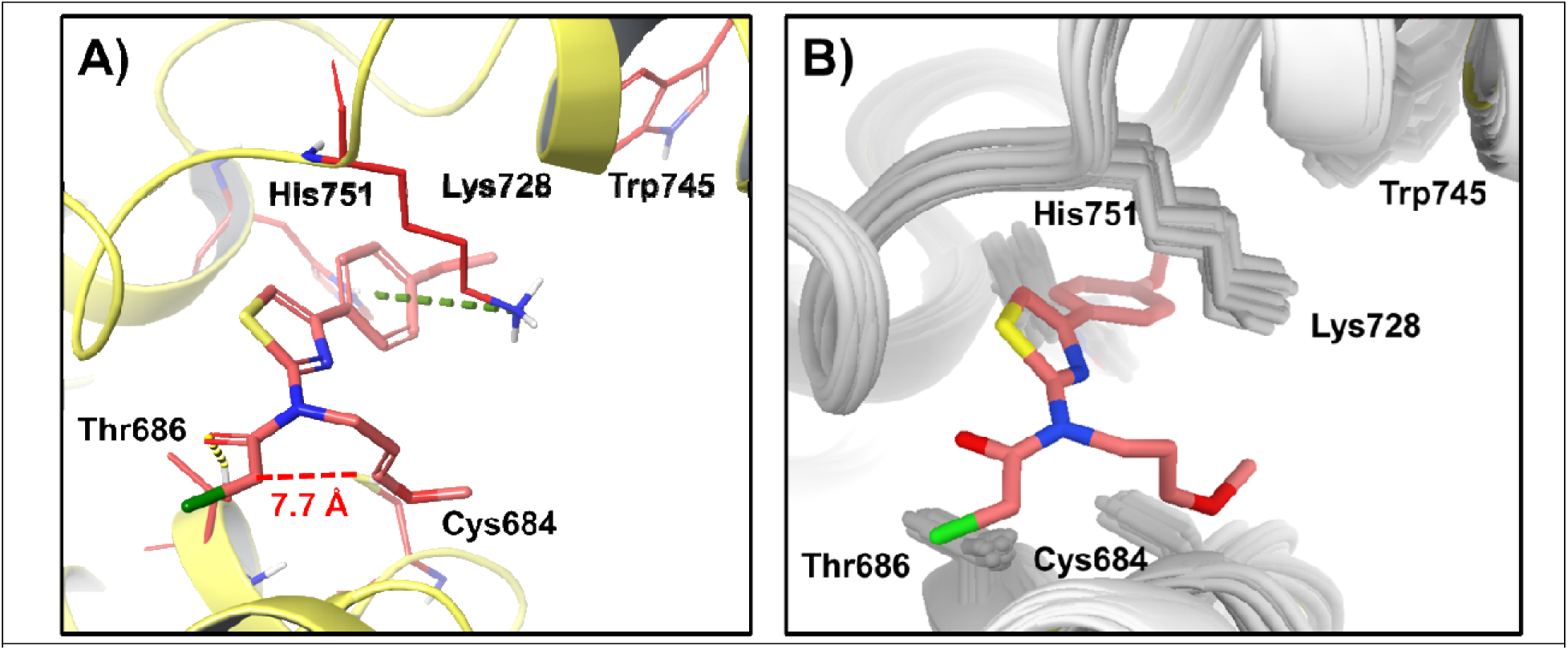
A) Predicted JT010-AaTRPA1 interaction. The distance between Cys684 and JT010 electrophilic site is 7.7 Å. JT010 (pink) forms a cation-π interaction (green line) with Lys728, a π-π stacking with His751, and a hydrogen bond (yellow line) with Thr686. B) Structural alignment of apo AaTRPA1 (gray) with the AaTRPA1-JT010 complex (yellow and pink).

## 4. Discussion

The results of the present study refuted our hypothesis that CDIAL was a more potent agonist of AaTRPA1 vs. HsTRPA1. Instead, CDIAL showed similar agonistic potencies of approximately 2 µM for both channels, suggesting CDIAL is a broad TRPA1 agonist. The agonistic potency of CDIAL for HsTRPA1 is within the range of 0.4-8.1 µM previously reported for other drimane sesquiterpene dialdehydes (i.e., polygodial and warburganal) against mammalian TRPA1 channels (Escalera et al., 2008; Mathie et al., 2017). Our findings with AaTRPA1 are consistent with our previous study that demonstrated 1) 10 µM CDIAL agonized AgTRPA1 expressed heterologously in *Xenopus* oocytes and 2) CDIAL elicited weak antifeedant responses in TRPA1-deficient *Ae. aegypti* (Inocente et al., 2018). Our results are the first to demonstrate that CDIAL is an agonist of HsTRPA1. This finding may explain the antinociceptive effects attributed to *Cinnamosma* bark (which is highly enriched with CDIAL) used in traditional medicines (Frimpong et al., 2021; Randrianarivelojosia et al., 2003). For example, other TRPA1 agonists, such acetaminophen and cannabinoids, are known to relieve pain via TRPA1-mediated inhibition of neuron excitability and neurotransmitter release (Andersson et al., 2011).

The agonistic potency and efficacy of CDIAL against both AaTRPA1 and HsTRPA1 were superior compared to that of nepetalactone. However, nepetalactone showed superior specificity for AaTRPA1 over HsTRPA1 compared to CDIAL. Our results are consistent with those of Melo et al. (Melo et al., 2021) who found that 1) catnip oil, which was presumed to contain 80-90% nepetalactone, was a more potent agonist of AaTRPA1 than HsTRPA1, and 2) 100 µM nepetalactone consistently agonized AaTRPA1. The potency of nepetalactone for AaTRPA1 estimated in the present study (197.0 μM) is within the range of potencies recently found for nepetalactone against 3 different hemipteran insect TRPA1 channels (172.5-698.4 μM) (Li et al., 2025). The high specificity of nepetalactone for insect TRPA1 over HsTRPA1 is attributed to the highly conserved ‘CVT’ motif present in the coupling domains of insect, but not mammalian, TRPA1 channels (i.e., Cys^684^VT in Ag/AaTRPA1, Fig S1) (Li et al., 2025).

CDIAL was a superior agonist of AaTRPA1 compared to JT010; the latter did not detectably agonize AaTRPA1 up to 10 µM. However, JT010 was a superior agonist of HsTRPA1 compared to CDIAL (∼3-fold more potent). The high selectivity of JT010 for HsTRPA1 over AaTRPA1 is not surprising given that JT010 is remarkably selective for HsTRPA1 over mouse TRPA1 (Matsubara et al., 2022; Takaya et al., 2015). The high affinity of JT010 for HsTRPA1 is attributed to covalent binding with Cys621 (analogous to Cys684 in AaTRPA1), which requires coordination by Phe669 (Matsubara et al., 2022; Takaya et al., 2015). While Cys621 is conserved among mosquito, human, and mouse TRPA1, Phe669 is only found in HsTRPA1 (Fig. S1). Our computational modeling results suggest that JT010 can occupy the conserved active site in AaTRPA1; however, differences in binding pocket composition result in an altered binding pose. Consequently, JT010 cannot form a covalent bond with Cys684 because its electrophilic site remains too distant, explaining why JT010 cannot induce the conformational changes necessary for channel activation in AaTRPA1.

The mutagenesis experiments with AaTRPA1 supported our hypothesis that the previously identified Lys residues (Lys656, Lys678, Lys728, Lys738, and Lys744) and/or Cys684 in the coupling domain of the NH_2_-terminus contributed to the binding of CDIAL. In particular, the Lys residues appear to play the major role in the binding of CDIAL compared to Cys684 given that CDIAL activation of the Cys684Ser mutant was similar to WT AaTRPA1, whereas activation of the Lys_null mutant was dramatically reduced compared to WT AaTRPA1. This finding is consistent with Mathie et al (Mathie et al., 2017) who demonstrated that dialdehyde sesquiterpenes structurally similar to CDIAL (i.e., polygodial and warburganal) were more reactive towards lysine compared to cysteine. It is important to note that mutation of all six Lys residues (with or without Cys684) did not completely abolish CDIAL agonism. A highly diminished, but detectable, activation by CDIAL was still observed in both the Lys_null and Cys684Ser + Lys_null constructs. Thus, it is likely that other residue(s) within and/or outside of the coupling domain provide minor contributions to CDIAL binding and activation.

Consistent with this notion, our molecular docking simulations not only highlighted Lys681 of AaTRPA1 (one of the Lys residues mutated in the Lys_null construct) as a residue of predicted importance in binding to CDIAL, but also revealed potential roles for Tyr651, Tyr654, and Asp680 in stabilizing the interaction. Notably, CDIAL binding appeared to induce a local conformational shift within the loop spanning Lys678 to Pro683, which includes Lys681. Future mutagenesis studies testing the role of each Lys residue that was mutated in the AaTRPA1 Lys_null construct, and assessing the putative roles of Tyr651, Tyr654, and Asp680 will be required to further delineate both the binding site(s) and mechanism(s) of CDIAL to AaTRPA1.

Two of the above Lys residues in the Lys_null mutant of AaTRPA1 (Lys656 and Lys738) are conserved in HsTRPA1 (Fig. S1), but neither were found to interact with CDIAL in the molecular docking simulations of AaTRPA1 or HsTRPA1. Thus, CDIAL likely interacts with AaTRPA1 and HsTRPA1 via independent mechanisms. In HsTRPA1, molecular docking simulations predicted a major role of Cys621 in the binding of CDIAL, a residue known to be involved with binding of JT010 and other electrophiles to HsTRPA1 (Matsubara et al., 2022; Takaya et al., 2015). Notably, the docking simulations also predicted that binding of CDIAL to Cys621 of HsTRPA1 induced a conformational shift of Pro666 similar to that which occurs for JT010 (Matsubara et al., 2022). Thus, CDIAL potentially binds to HsTRPA1 via a mechanism similar to JT010. Future mutagenesis studies will be required to demonstrate that Cys621 and Pro666 contribute to the binding of CDIAL to HsTRPA1.

In summary, our results provide an important advance in our understanding of how drimane sesquiterpenes interact with TRPA1 channels, which to date has not been well characterized. We identified a major binding pocket for CDIAL in the coupling domain of AaTRPA1 that involves one or more Lys residues. Although the amino acid-sequence novelty of this binding pocket was not well conserved in HsTRPA1, CDIAL was a highly potent agonist of HsTRPA1. Modeling results suggest that CDIAL binds to HsTRPA1 through a distinct mechanism that more closely resembles that of JT010. Further characterization of species-specific binding mechanisms of CDIAL to TRPA1 channels can potentially be used for designing or discovering highly potent, mosquito-specific agonists of TRPA1 with use as biorational repellents or highly potent agonists of HsTRPA1 with use as drugs that mimic the therapeutic effects of some traditional medicines.

## Acknowledgments

The authors thank Drs. Shigeru Saito and Makoto Tominaga (National Institute for Physiological Science, Okazaki, Japan) for providing the HsTRPA1 cDNA construct. The present study was supported by a grant from the National Institute of Allergy and Infectious Diseases of the National Institutes of Health (1R56AI158674-01A1), The Ohio State University (OSU) College of Pharmacy Dean’s Innovation Award, and State and Federal Funds Appropriated to the OSU College of Food, Agricultural, and Environmental Sciences Wooster Campus.

## Appendix A. Supplementary data

**Fig. S1.**
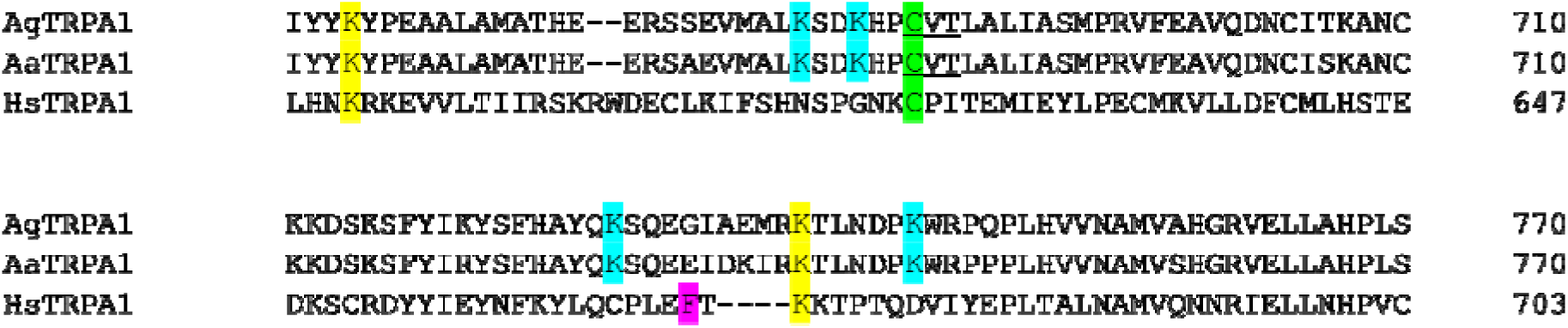
Amino acid sequence alignment of the coupling domain of mosquito (Ag, Aa) and human (Hs) TRPA1 channels. Green-shaded residues correspond to a conserved Cys (683 in Ag/AaTRPA1 or 621 in HsTRPA1) predicted to be important in the covalent binding of electrophiles, including JT010 and CDIAL. Yellow-shaded residues indicate conserved Lys residues in both mosquito and human TRPA1 (656 and 738 in Ag/AaTRPA1 or 591and 671 in HsTRPA1) predicted to be important in the binding of CDIAL. Blue-shaded residues indicate Lys residues only conserved in mosquito TRPA1 (678, 681, 728, 744 in Ag/AaTRPA1) predicted to be important in binding CDIAL. Magenta-shaded residue is Phe669 that together with Cys621 contributes to binding of JT010 to HsTRPA1(Matsubara et al., 2022). Underlined residues indicate conserved ‘CVT’ motif of insect TRPA1 channels involved with binding of nepetalactone (Li et al., 2025). For mutated constructs of AaTRPA1 used in the present study the highlighted Cys was changed to Ser and/or all of the highlighted Lys residues were simultaneously changed to Ala.

**Fig. S2.**
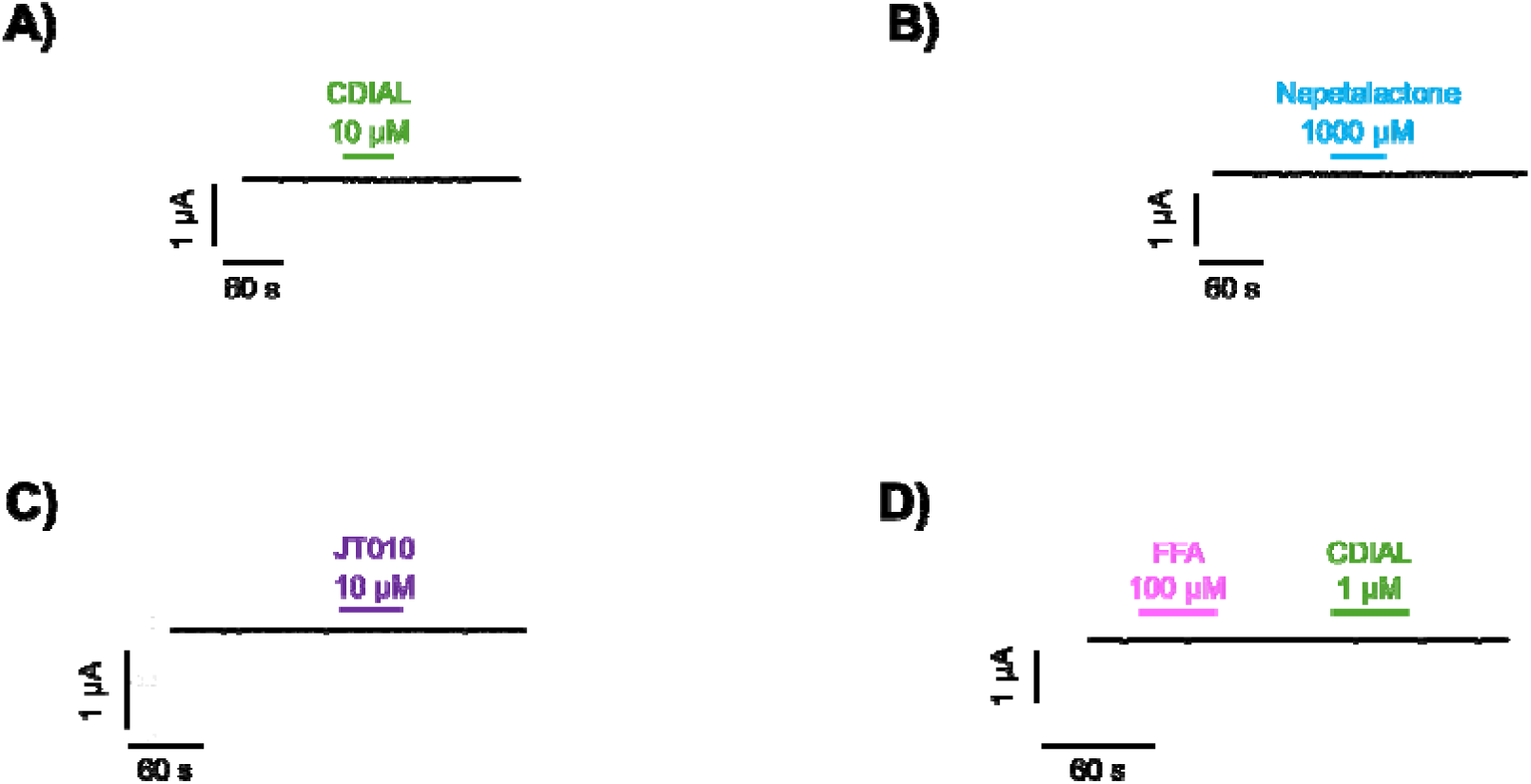
Representative traces of I_m_ in voltage-clamped, H_2_O-injected oocytes in response to CDIAL (A), nepetalactone (B), JT010 (C), or FFA and CDIAL (D). In panels A-C, only the highest concentration used in the corresponding concentration-response experiment is shown. In all cases, the ΔI_m_ responses were below detectable limits.

**Fig. S3.**
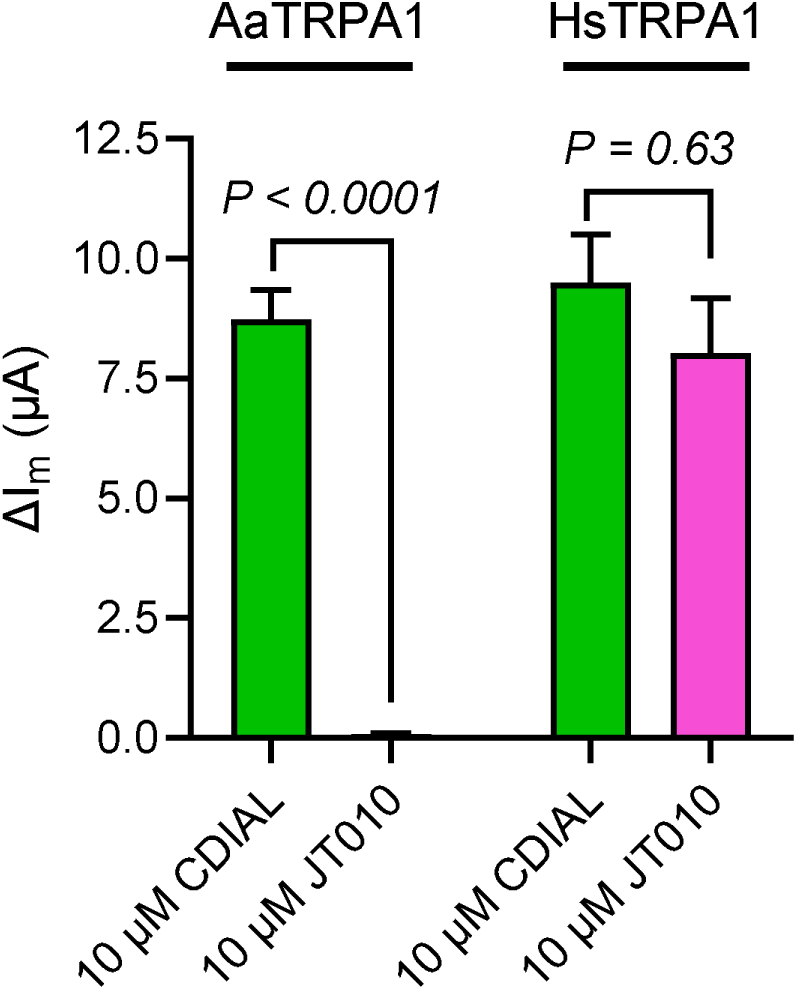
Comparisons of the agonistic responses induced by 10 µM CDIAL and 10 µM JT010 in AaTRPA1 and HsTRPA1 oocytes. Values are means ± SEM, based on independent oocytes from the same batch (in AaTRPA1, n = 3 for CDIAL and n = 5 for JT010; in HsTRPA1, n = 3 for CDIAL and n = 4 for JT010). For AaTRPA1, *P* value is derived from an unpaired t-test in Prism for AaTRPA1 (data were normally distributed as determined by a Shapiro-Wilk test). For HsTRPA1, *P* value is derived from a Mann-Whitney test in Prism (data were not normally distributed as determined by a Shapiro-Wilk test).

**Fig. S4.**
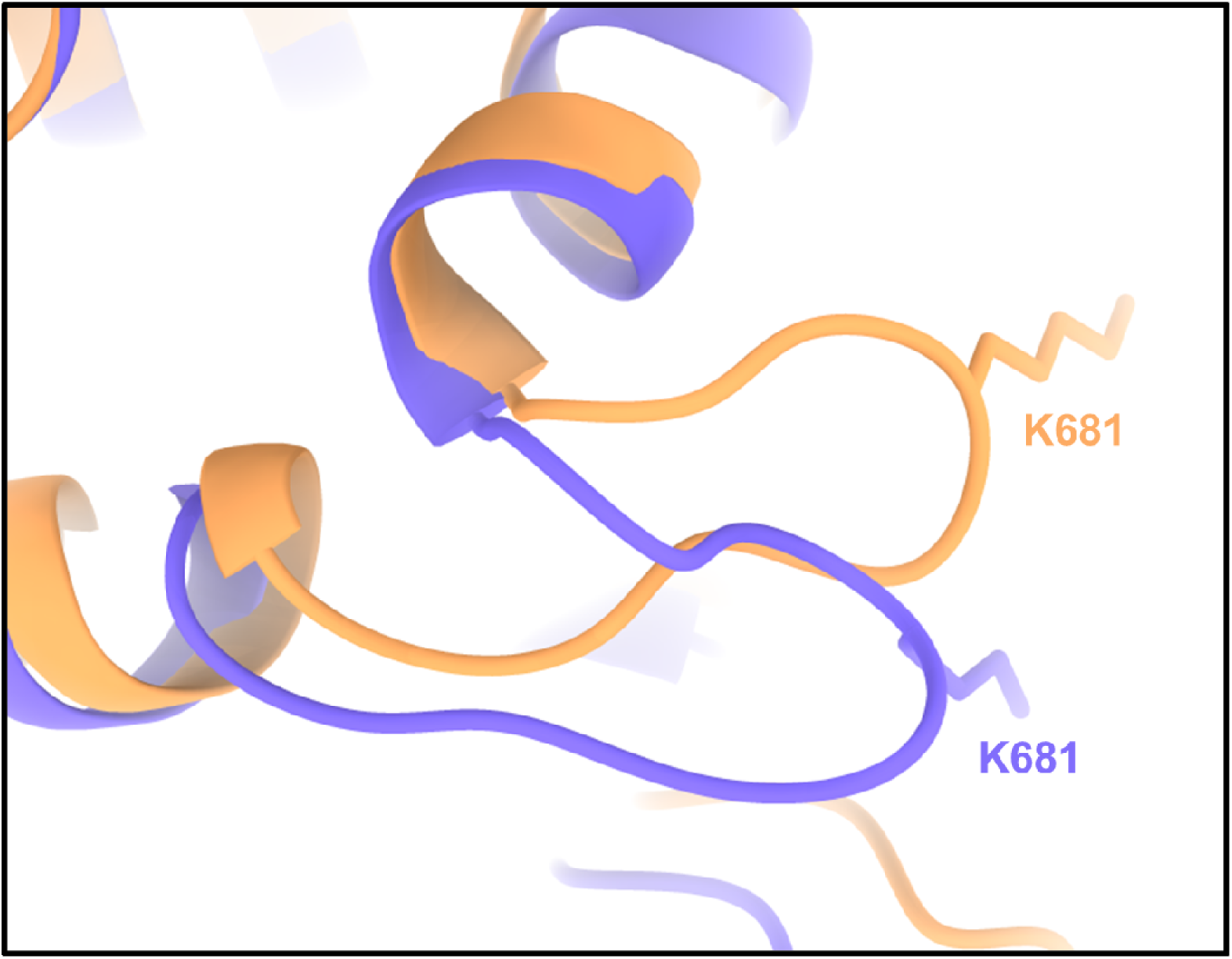
Local conformational shift induced by covalent binding of CDIAL. Orange represents the AF3 predicted apo AaTRPA1 structure, while the purple represents the AaTRPA1-CDIAL covalent complex.

## Notes

### Competing Interest Statement

The authors have declared no competing interest.

## References

Abramson, J., Adler, J., Dunger, J., Evans, R., Green, T., Pritzel, A., Ronneberger, O., Willmore, L., Ballard, A.J., Bambrick, J., Bodenstein, S.W., Evans, D.A., Hung, C.C., O’Neill, M., Reiman, D., Tunyasuvunakool, K., Wu, Z., Zemgulyte, A., Arvaniti, E., Beattie, C., Bertolli, O., Bridgland, A., Cherepanov, A., Congreve, M., Cowen-Rivers, A.I., Cowie, A., Figurnov, M., Fuchs, F.B., Gladman, H., Jain, R., Khan, Y.A., Low, C.M.R., Perlin, K., Potapenko, A., Savy, P., Singh, S., Stecula, A., Thillaisundaram, A., Tong, C., Yakneen, S., Zhong, E.D., Zielinski, M., Zidek, A., Bapst, V., Kohli, P., Jaderberg, M., Hassabis, D., Jumper, J.M., 2024. Accurate structure prediction of biomolecular interactions with AlphaFold 3. Nature 630, 493–500.

Andersson, D.A., Gentry, C., Alenmyr, L., Killander, D., Lewis, S.E., Andersson, A., Bucher, B., Galzi, J.-L., Sterner, O., Bevan, S., Högestätt, E.D., Zygmunt, P.M., 2011. TRPA1 mediates spinal antinociception induced by acetaminophen and the cannabinoid Δ9-tetrahydrocannabiorcol. Nature Communications 2, 551.

Chandel, A., DeBeaubien, N.A., Ganguly, A., Meyerhof, G.T., Krumholz, A.A., Liu, J., Salgado, V.L., Montell, C., 2024. Thermal infrared directs host-seeking behaviour in Aedes aegypti mosquitoes. Nature 633, 615–623.

Corfas, R.A., Vosshall, L.B., 2015. The cation channel TRPA1 tunes mosquito thermotaxis to host temperatures. eLife 4, e11750.

Escalera, J., von Hehn, C.A., Bessac, B.F., Sivula, M., Jordt, S.E., 2008. TRPA1 mediates the noxious effects of natural sesquiterpene deterrents. J Biol Chem 283, 24136–24144.

Frimpong, E.K., Asong, J.A., Aremu, A.O., 2021. A Review on Medicinal Plants Used in the Management of Headache in Africa. Plants 10, 2038.

Gracheva, E.O., Ingolia, N.T., Kelly, Y.M., Cordero-Morales, J.F., Hollopeter, G., Chesler, A.T., Sánchez, E.E., Perez, J.C., Weissman, J.S., Julius, D., 2010. Molecular basis of infrared detection by snakes. Nature 464, 1006–1011.

Gupta, R., Saito, S., Mori, Y., Itoh, S.G., Okumura, H., Tominaga, M., 2016. Structural basis of TRPA1 inhibition by HC-030031 utilizing species-specific differences. Sci Rep 6, 37460.

Hinman, A., Chuang, H.H., Bautista, D.M., Julius, D., 2006. TRP channel activation by reversible covalent modification. Proc Natl Acad Sci USA 103, 19564–19568.

Hu, H., Bandell, M., Petrus, M.J., Zhu, M.X., Patapoutian, A., 2009. Zinc activates damage-sensing TRPA1 ion channels. Nat Chem Biol 5, 183–190.

Hu, H., Tian, J., Zhu, Y., Wang, C., Xiao, R., Herz, J.M., Wood, J.D., Zhu, M.X., 2010. Activation of TRPA1 channels by fenamate nonsteroidal anti-inflammatory drugs. Pflügers Archiv - European Journal of Physiology 459, 579–592.

Inocente, E.A., Nguyen, B., Manwill, P.K., Benatrehina, A., Kweka, E., Wu, S., Cheng, X., Rakotondraibe, L.H., Piermarini, P.M., 2019. Insecticidal and antifeedant activities of Malagasy medicinal plant (*Cinnamosma* sp.) extracts and drimane-type sesquiterpenes against *Aedes aegypti m*osquitoes. Insects 10, 373.

Inocente, E.A., Shaya, M., Acosta, N., Rakotondraibe, L.H., Piermarini, P.M., 2018. A natural agonist of mosquito TRPA1 from the medicinal plant *Cinnamosma fragrans* that is toxic, antifeedant, and repellent to the yellow fever mosquito *Aedes aegypti*. PLoS Negl Trop Dis 12, e0006265.

Kang, K., Pulver, S.R., Panzano, V.C., Chang, E.C., Griffith, L.C., Theobald, D.L., Garrity, P.A., 2010. Analysis of *Drosophila* TRPA1 reveals an ancient origin for human chemical nociception. Nature 464, 597–600.

Kwon, Y., Kim, S.H., Ronderos, D.S., Lee, Y., Akitake, B., Woodward, O.M., Guggino, W.B., Smith, D.P., Montell, C., 2010. *Drosophila* TRPA1 channel is required to avoid the naturally occurring insect repellent citronellal. Curr Biol 20, 1672–1678.

Li, J., Wang, B., Wang, Y., Li, F., Li, Z., Liu, X., Zhang, S., 2025. TRPA1-kinase axis polarization: Nepetalactone drives pest repulsion and predator attraction via divergent PKC/CaMKII signaling. Journal of Advanced Research.

Li, T., Saito, C.T., Hikitsuchi, T., Inoguchi, Y., Mitsuishi, H., Saito, S., Tominaga, M., 2019. Diverse sensitivities of TRPA1 from different mosquito species to thermal and chemical stimuli. Sci Rep 9, 20200.

Lv, H., Wang, G., Wu, X., Jiang, D., 2025. Molecular cloning and functional characterizations of transient receptor potential A1 (TRPA1) in Aedes albopictus. Pesticide Biochemistry and Physiology 208, 106295.

Macpherson, L.J., Dubin, A.E., Evans, M.J., Marr, F., Schultz, P.G., Cravatt, B.F., Patapoutian, A., 2007. Noxious compounds activate TRPA1 ion channels through covalent modification of cysteines. Nature 445, 541–545.

Maekawa, E., Aonuma, H., Nelson, B., Yoshimura, A., Tokunaga, F., Fukumoto, S., Kanuka, H., 2011. The role of proboscis of the malaria vector mosquito *Anopheles stephensi* in host-seeking behavior. Parasites & Vectors 4, 10.

Manwill, P.K., Kalsi, M., Wu, S., Martinez Rodriguez, E.J., Cheng, X., Piermarini, P.M., Rakotondraibe, H.L., 2020. Semi-synthetic cinnamodial analogues: Structural insights into the insecticidal and antifeedant activities of drimane sesquiterpenes against the mosquito *Aedes aegypti*. PLoS Neglected Tropical Diseases 14, e0008073.

Mathie, K., Lainer, J., Spreng, S., Dawid, C., Andersson, D.A., Bevan, S., Hofmann, T., 2017. Structure–pungency relationships and TRP channel activation of drimane sesquiterpenes in Tasmanian Pepper (*Tasmannia lanceolata*). J Agric Food Chem 65, 5700–5712.

Matsubara, M., Muraki, Y., Hatano, N., Suzuki, H., Muraki, K., 2022. Potent Activation of Human but Not Mouse TRPA1 by JT010. International Journal of Molecular Sciences 23, 14297.

Melo, N., Capek, M., Arenas, O.M., Afify, A., Yilmaz, A., Potter, C.J., Laminette, P.J., Para, A., Gallio, M., Stensmyr, M.C., 2021. The irritant receptor TRPA1 mediates the mosquito repellent effect of catnip. Curr Biol 31, 1988–1994 e1985.

Mitchell, R.D., Zhu, J., Carr, A.L., Dhammi, A., Cave, G., Sonenshine, D.E., Roe, R.M., 2017. Infrared light detection by the haller’s organ of adult american dog ticks, Dermacentor variabilis (Ixodida: Ixodidae). Ticks and Tick-borne Diseases 8, 764–771.

Park, Y., Piermarini, P.M., 2025. Heat activation desensitizes Aedes aegypti transient receptor potential ankyrin 1 (AaTRPA1) to chemical agonists that repel mosquitoes. Pesticide Biochemistry and Physiology 209, 106326.

Randrianarivelojosia, M., Rasidimanana, V.T., Rabarison, H., Cheplogoi, P.K., Ratsimbason, M., Mulholland, D.A., Mauclère, P., 2003. Plants traditionally prescribed to treat tazo (malaria) in the eastern region of Madagascar. Malaria Journal 2, 25.

Romero, M.F., Fong, P., Berger, U.V., Hediger, M.A., Boron, W.F., 1998. Cloning and functional expression of rNBC, an electrogenic Na+-HCO3- cotransporter from rat kidney. Am J Physiol 274, F425–432.

Salgado, V.L., 2017. Insect TRP channels as targets for insecticides and repellents. J Pest Sci 42, 1–6.

Sastry, G.M., Adzhigirey, M., Day, T., Annabhimoju, R., Sherman, W., 2013. Protein and ligand preparation: parameters, protocols, and influence on virtual screening enrichments. J Comput Aided Mol Des 27, 221–234.

Schrödinger, Inc., 2025. Schrödinger Release 2025-3. Maestro, Schrödinger, LLC, New York, NY.

Takaya, J., Mio, K., Shiraishi, T., Kurokawa, T., Otsuka, S., Mori, Y., Uesugi, M., 2015. A Potent and Site-Selective Agonist of TRPA1. Journal of the American Chemical Society 137, 15859–15864.

Talavera, K., Startek, J.B., Alvarez-Collazo, J., Boonen, B., Alpizar, Y.A., Sanchez, A., Naert, R., Nilius, B., 2020. Mammalian Transient Receptor Potential TRPA1 Channels: From Structure to Disease. Physiological Reviews 100, 725–803.

Wang, G., Qiu, Y.T., Lu, T., Kwon, H.W., Pitts, R.J., Van Loon, J.J., Takken, W., Zwiebel, L.J., 2009. *Anopheles gambiae* TRPA1 is a heat-activated channel expressed in thermosensitive sensilla of female antennae. Eur J Neurosci 30, 967–974.

Wang, S., Lee, J., Ro, J.Y., Chung, M.-K., 2012. Warmth suppresses and desensitizes damage-sensing ion channel TRPA1. Molecular Pain 8, 22.

Zhang, M., Ma, Y., Ye, X., Zhang, N., Pan, L., Wang, B., 2023. TRP (transient receptor potential) ion channel family: structures, biological functions and therapeutic interventions for diseases. Signal Transduction and Targeted Therapy 8, 261.

